# Cool temperature limit Zika virus genome replication

**DOI:** 10.1101/2021.06.07.447476

**Authors:** Blanka Tesla, Jenna S. Powers, Yvonne Barnes, Shamil Lakhani, Marissa D. Acciani, Courtney C. Murdock, Melinda A. Brindley

## Abstract

Zika virus is a mosquito-borne flavivirus known to cause severe birth defects and neuroimmunological disorders. We have previously demonstrated that mosquito transmission of Zika virus decreases with temperature. While transmission was optimized at 29°C, it was limited at cool temperatures (< 22°C) due to poor virus establishment in the mosquitoes. Temperature is one of the strongest drivers of vector-borne disease transmission due to its profound effect on ectothermic mosquito vectors, viruses, and their interaction. Although there is substantial evidence of temperature effects on arbovirus replication and dissemination inside mosquitoes, little is known about whether temperature affects virus replication directly or indirectly through mosquito physiology. In order to determine the mechanisms behind temperature-induced changes in Zika virus transmission potential, we investigated different steps of the virus replication cycle in mosquito cells (C6/36) at optimal (28°C) and cool (20°C) temperatures. We found that cool temperature did not alter Zika virus entry or translation but reduced the amount of double-stranded RNA replication intermediates. If replication complexes were first formed at 28°C and the cells were subsequently shifted to 20°C, the late steps in the virus replication cycle were efficiently completed. These data suggest that cool temperature decreases the efficiency of Zika virus genome replication in mosquito cells. This phenotype was observed in the Asian-lineage of Zika virus, while the African-lineage Zika virus was less restrictive at 20°C.

**Importance:** With half of the human population at risk, arboviral diseases represent a substantial global health burden. Zika virus, previously known to cause sporadic infections in humans, emerged in the Americas in 2015 and quickly spread worldwide. There was an urgent need to better understand the disease pathogenesis, develop therapeutics and vaccines, as well as to understand, predict, and control virus transmission. In order to efficiently predict the seasonality and geography for Zika virus transmission, we need a deeper understanding of the host-pathogen interactions and how they can be altered by environmental factors such as temperature. Identifying the step in the virus replication cycle that is inhibited in cool conditions can have implications in modeling the temperature suitability for arbovirus transmission as global environmental patterns change. Understanding the link between pathogen replication and environmental conditions can potentially be exploited to develop new vector control strategies in the future.

## Introduction

Diseases such as Zika, dengue, and chikungunya, once considered tropical and sub-tropical diseases, have spread explosively throughout the world due to climate change, globalization, and increased urbanization. With half of the human population at risk, arboviral diseases represent a substantial global health burden (1). Zika virus (ZIKV) is a mosquito-borne flavivirus known to cause sporadic and mild infections in humans. In 2015, ZIKV emerged in the Americas and within a year quickly spread to approximately 65 countries worldwide, resulting in over 360,000 suspected cases (2). Shortly after it was linked to birth defects (3) and neuroimmunological disorders (Guillain–Barré syndrome) (4), ZIKV was declared a “public health emergency of international concern” (5). With no therapeutics or vaccines to mitigate disease, ZIKV was quickly put at the forefront of research interest to fulfill the urgent need to better understand disease transmission, pathogenesis, and prevention.

Temperature is one of the strongest drivers of vector-borne disease transmission due to its profound impact on ectothermic mosquito vectors, pathogens, and their interactions (6). Numerous studies have investigated the effects of temperature on mosquito infection, the extrinsic incubation period (EIP), and overall transmission potential in a diversity of vector-borne disease systems (7–11). However, the exact mechanisms of how temperature shapes arboviral transmission are rarely elucidated. Temperature can either affect virus replication directly, or indirectly by altering mosquito physiology (12), immunity (13, 14), and cellular processes (15). Even though arboviruses evolved to replicate across a wide range of temperatures, from within invertebrate vectors to febrile mammalian and avian hosts, temperature can also modify virus replication. Studies have shown that temperature can alter virus structure (16, 17), induce the fluctuation of viral surface proteins required for entry in the host cell (18, 19), or affect genome replication (20, 21). Although a few studies have investigated ZIKV structure and thermal stability (22–24), no studies have investigated how temperature affects the ZIKV replication cycle.

In our previous work, we demonstrated that ZIKV transmission in *Aedes aegypti* mosquitoes was optimized at 29°C and had a thermal range of 22.7°C to 34.7°C. Although warm temperatures facilitated rapid virus replication, warm conditions (>36°C) also increased mosquito mortality and led to an overall decrease in transmission potential. However, cool conditions (<22°C) prevented the mosquitoes from becoming infectious due to poor midgut infection and escape (25). In order to understand the mechanisms of reduced ZIKV transmission potential at suboptimal temperatures, we investigated ZIKV replication *in vitro* using a mosquito cell line incubated across a range of temperatures. Here we show ZIKV replication in cell culture closely mirrors the findings in mosquitoes. ZIKV replication occurs faster as the temperature increases, until the temperature starts to induce cell death, while cool temperatures significantly decrease the replication kinetics. We formulated two, not mutually exclusive, hypotheses to address our observations: 1) The stress from cool temperatures alters the cellular environment, either limiting host factors necessary for viral replication or overproducing an inhibitory factor not produced at permissive temperatures. 2) The suboptimal temperatures prevent a viral function required to complete the viral replication cycle and produce progeny virus. We found that reduced ZIKV replication at suboptimal temperature is not a result of acute cellular stress. In addition, maintaining cells at 20°C did not appear to affect early and late steps in the viral replication cycle; however, it did affect genome replication. While American isolates of ZIKV were restricted, African isolates were able to replicate more efficiently at 20°C.

## Results

### ZIKV, DENV, and CHIKV replication curves at different temperatures

To characterize the effect of temperature on ZIKV replication in mosquito cells, we infected C6/36 cells with ZIKV and incubated them at six temperatures ranging from 16°C to 36°C (Fig 1A). To mirror our experiment in mosquitoes (25), six independent plates were infected at 28°C for two hours, inoculum was removed, and the plates were then moved to the indicated temperatures where they were maintained for the remainder of the experiment. Initial ZIKV replication and peak titers occurred more quickly as the temperature increased. Incubation at 16°C resulted in almost complete inhibition of virus production in C6/36 cells, while incubation at 20°C resulted in delayed particle release and more than a 5-log reduction in peak titers. While ZIKV replication started robustly at 36°C, virus replication was drastically reduced over time. This phenotype is a result of temperature-related cellular stress and was not observed in ZIKV-infected mammalian cells that optimally produce virus at 37°C (Fig 1B). While virus replication peaked at 2.67 × 10^9^ TCID_50_/mL at 28°C in mosquito cells, the peak titer was 3.5-log lower at 28°C in Vero cells. In comparison, Vero cells infected at 37°C produced the highest titers of virus throughout the experiment, yet titers in C6/36 cells at 36°C peaked low and early and then fell over the course of the experiment. This suggests that peak virus replication is determined by both intracellular components and optimal environmental conditions.

**FIG 1.**
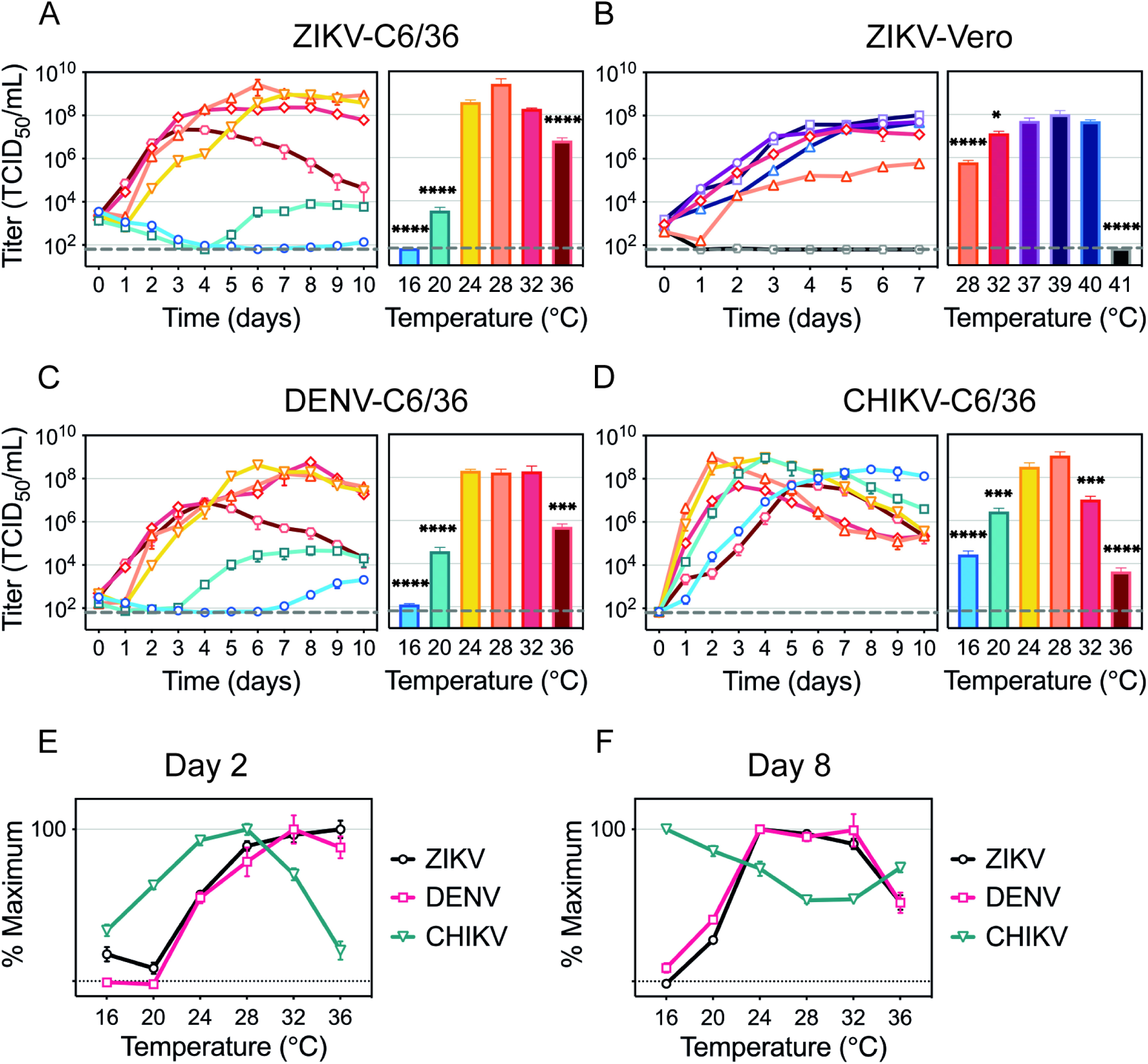
Temperature effects on ZIKV, DENV, and CHIKV replication. Replication rates for (**A**) ZIKV in C6/36 cells and titers on day 6, (**B**) ZIKV in Vero cells and titers on day 7, (**C**) DENV in C6/36 cells and titers on day 7, and (**D**) CHIKV in C6/36 cells and titers on day 2. % maximum on (**E**) day 2 and (**F**) day 8. C6/36 cells were incubated with ZIKV (MOI 0.1), DENV (MOI 0.1), or CHIKV (MOI 0.005) for 2 h at 28°C, when infectious media was removed and replaced with fresh media. The cells were kept at 16°C, 20°C, 24°C, 28°C, 32°C, or 36°C for 10 d. Vero cells were infected with ZIKV (MOI 0.1) for 1 h at 37°C. The cells were incubated at 28°C, 32°C, 37°C, 39°C, 40°C, or 41°C for 7 d. Supernatants were collected every 24 h and titrated on Vero cells. Titers are displayed in TCID_50_ units/mL. Dashed lines represent the limit of detection. Data shown are means ± SEM from three independent replicates for each virus. Bar graphs represent viral titers on the day peak titers were reached at optimal temperature (28°C for C6/36, 37°C for Vero). Statistical differences were determined with a one-way ANOVA on log transformed data for each experiment. Dunnett’s test was used for multiple comparisons, where optimal temperature (28°C or 37°C) was compared to all other temperatures. * *p* < 0.05, *** *p* < 0.001, **** *p* < 0.0001

Next, we compared ZIKV replication across six temperatures to another flavivirus (DENV) and to an alphavirus (CHIKV). DENV had similar replication dynamics to ZIKV, yet DENV was less restricted at cool temperatures, reaching 1.45 × 10^3^ TCID_50_/mL at 16°C and 3.74 × 10^6^ TCID_50_/mL at 20°C (Fig 1C). CHIKV replication kinetics and its response to temperature greatly differed relative to both flaviviruses (Fig 1D). In order to compare how the temperature differentially affects the three viruses, we compared how much virus was produced across the temperature range early (Fig. 1E) and late (Fig. 1F) during the infection. While CHIKV replication proceeded at a faster rate with both cool and warm temperatures inhibiting optimal viral yields early in the infection, ZIKV and DENV were observed to produce high titers as the temperature increased (Fig 1E). However, both ZIKV and DENV titers were low at 36°C late in infection, suggesting that prolonged high temperature treatment may enhance early replication, but eventually caused cell death (Fig. 1F). Interestingly, CHIKV peak TCID_50_ values were similar at 28°C and 20°C, simply delayed under cool conditions, suggesting C6/36 cells are capable of producing virus particles at 20°C; however, low temperatures might affect ZIKV and CHIKV replication differently.

### Temperature and virus effects on cell physiology

Temperature profoundly affects cell physiology and metabolism; therefore, it can alter cell proliferation, viability, and protein production. We monitored uninfected as well as ZIKV-, DENV-, and CHIKV-infected C6/36 cell proliferation and viability at temperatures ranging from 16°C to 36°C. The number of cell generations over a four-day period increased proportionally with temperature except for cells incubated at 36°C, where cell proliferation decreased (Fig 2A). While DENV infection did not alter cell proliferation at any temperature, ZIKV infection reduced the number of generations at 32°C and CHIKV decreased cell proliferation at all temperatures. Low and high temperatures also affected cell viability, represented by lower ATP levels, after cells were maintained at the indicated temperature for six days (Fig 2B). CHIKV was more cytopathic than ZIKV and DENV, while ZIKV affected cell viability only at warm temperatures. DENV-infected cells showed little-to-no reduction in ATP levels in comparison to uninfected cells.

**FIG 2.**
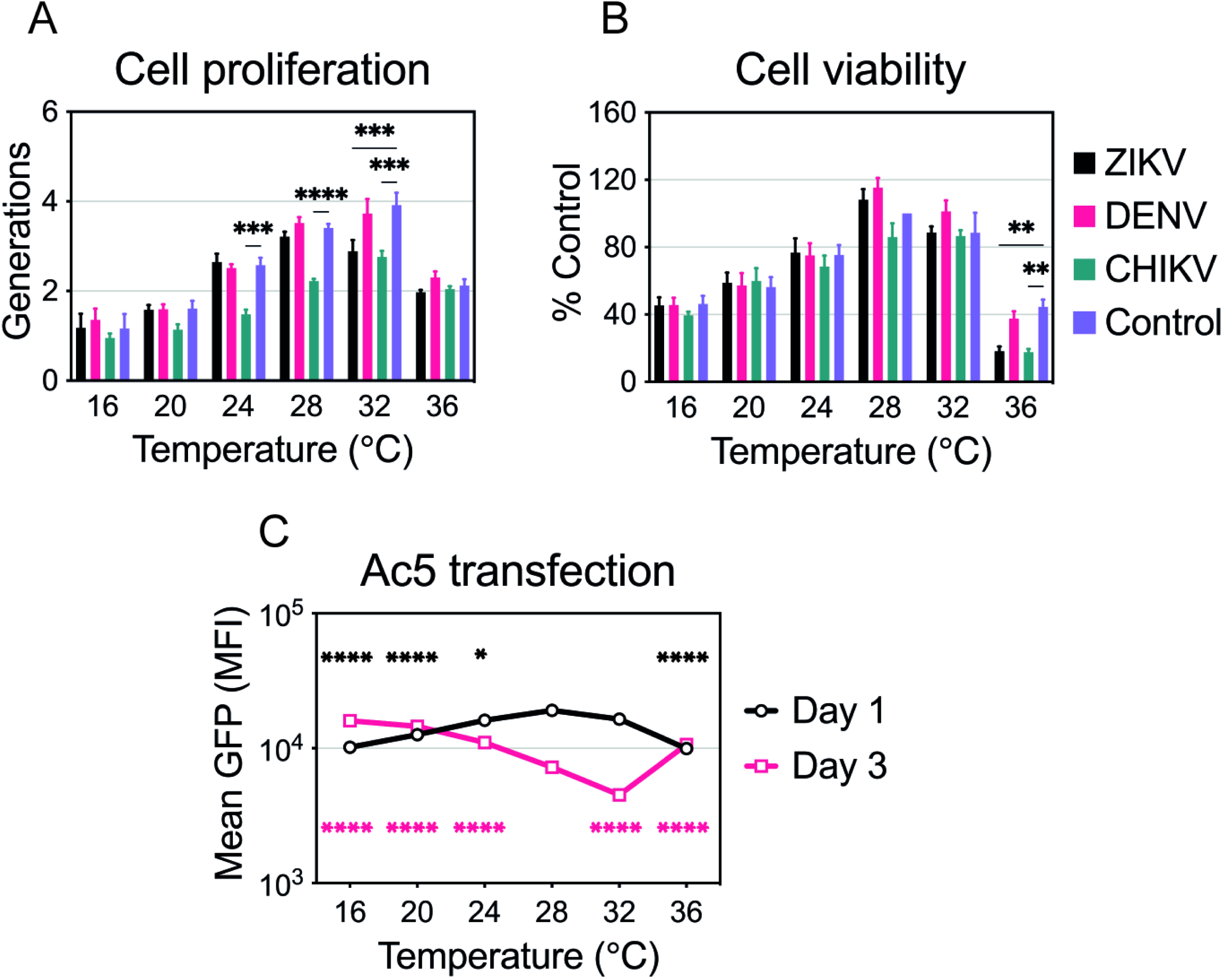
Temperature and virus effects on cell proliferation, viability, and protein production. (**A**) Cell proliferation of uninfected or infected C6/36 cells at different temperatures (16°C-36°C). Dyed cells were infected with ZIKV, DENV, or CHIKV as described in Fig 1 and were incubated at one of the six temperatures for 4 d. Intracellular fluorescence was normalized to uninfected cells on day 0. Data are presented as mean ± SEM from three independent experiments. (**B**) Cell viability of uninfected or infected C6/36 cells incubated at six temperatures (16°C-36°C) for 6 d. Cell viability was normalized to uninfected cells maintained at 28°C and data are presented as mean ± SEM from three independent experiments. Statistical differences for cell proliferation and viability were assessed by a two-way ANOVA with Dunnett’s correction for multiple comparisons. For each temperature, uninfected cells were compared to each virus. (**C**) Protein production at each temperature (16°C-36°C) on days 1 and 3 was assessed after transfecting C6/36 cells with Ac5-STABLE2-neo plasmid. The data shown, expressed as mean GFP, are means ± SEM from three replicates. Statistically significant mean fluorescence was assessed by using a two-way ANOVA on log transformed data followed by Dunnett’s test for multiple comparisons. For days 1 and 3, 28°C was compared to all other temperatures. The color of the significance symbol corresponds to the color for each day. * *p* < 0.05, ** *p* < 0.01, *** *p* < 0.001, **** *p* < 0.0001

In order to investigate the effects of temperature on protein production, we transfected C6/36 cells with an insect expression plasmid that encodes for GFP (pAc5-GFP) (Fig 2C). One day after transfection, cells produced less GFP at suboptimal (16°C, 20°C, 24°C) and hot (36°C) temperatures. Decreased protein production suggests that cells are slightly less metabolically active early after transfection at those temperatures. Three days after transfection, cells incubated at 28°C and 32°C were less fluorescent, likely due to faster cell proliferation. Slower proliferation at other temperatures resulted in higher mean GFP fluorescence as there were more GFP-positive cells. Therefore, while C6/36 cells do not proliferate as efficiently at low temperatures, they are still able to produce proteins.

### ZIKV, DENV, and CHIKV replication in 20°C-adapted C6/36 cells

When performing the replication curves shown in Figure 1, cells normally maintained at 28°C were infected and transferred to the different temperature treatments. Therefore, cells were subjected to temperature change during the initial infection time, which may have triggered acute cellular responses that would not be present if cells were maintained at the various temperature treatments prior to infection. To determine if virus replication was inhibited at 20°C due to acute cellular stress responses, we adapted C6/36 cells to grow at 20°C. After maintaining C6/36 cells at 20°C for several months, we compared the adapted cell morphology to normal cells and observed no changes. We also demonstrated that 20°C-adapted cells proliferated faster at 20°C than the cells that were shifted from 28°C to 20°C (Fig 3A and 3B). We then investigated ZIKV, DENV, and CHIKV replication in 20°C-adapted cells and found there was no difference in virus yields between adapted and non-adapted cells at 20°C (Fig 3C-E). This suggests that the reduced ZIKV replication was not likely a result of acute cellular stress brought on by the temperature shift.

**FIG 3.**
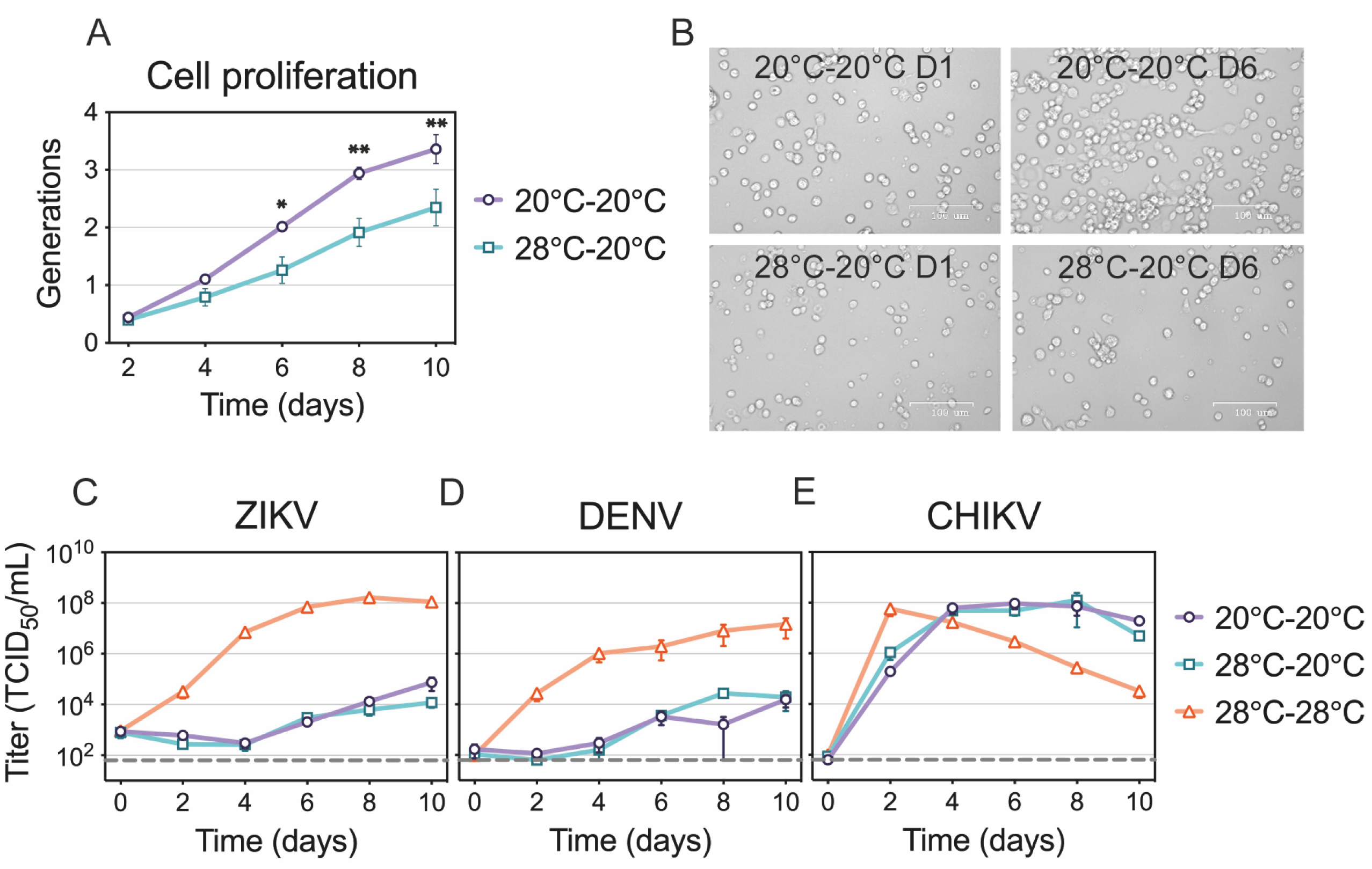
ZIKV, DENV, and CHIKV replication in 20°C-adapted C6/36 cells. (**A and B**) Cell proliferation of 20°C-adapted C6/36 cells in comparison to standard cells at 20°C. Both cell types were stained and maintained at 20°C for 10 d. Fluorescence was measured every 2 d and normalized to either 20°C-adapted or standard cells on day 0. Pictures of the cells were taken on days 0 and 6 and representative images are shown in panel B. The data shown are means ± SEM from three replicates. Statistical differences were determined by using a two-way ANOVA with Bonferroni’s correction where adapted cells were compared to standard cells at each time point. (**C**) Replication rates of ZIKV, (**D**) DENV, and (**E**) CHIKV in 20°C-adapted cells were compared to standard cells at 20°C and 28°C. 20°C-adapted cells were infected and kept at 20°C, while standard cells were infected at 28°C and kept at 20°C or 28°C. Supernatants were collected every 48 h and titrated. Dashed lines represent the limit of detection. Data shown are means ± SEM of results from three independent experiments for each virus. * *p* < 0.05, ** *p* < 0.01

### Effects of suboptimal temperature on ZIKV spread

We next sought out to determine which part of the virus replication cycle is affected by cool temperatures. We first examined if cells with established ZIKV infection are capable of producing infectious virus particles at 20°C (Fig 4A). Persistently infected C6/36 cells were distributed into two dishes, one of which was transferred to 20°C, while the other was kept at 28°C. Interestingly, the amount of infectious virus produced every 24 hours for five days was similar at 20°C and 28°C. We observed a slight decrease in virus yields, which was likely due to decreased cellular metabolism at 20°C. Overall, this suggests that later steps in virus replication are not inhibited at cool temperatures.

**FIG 4.**
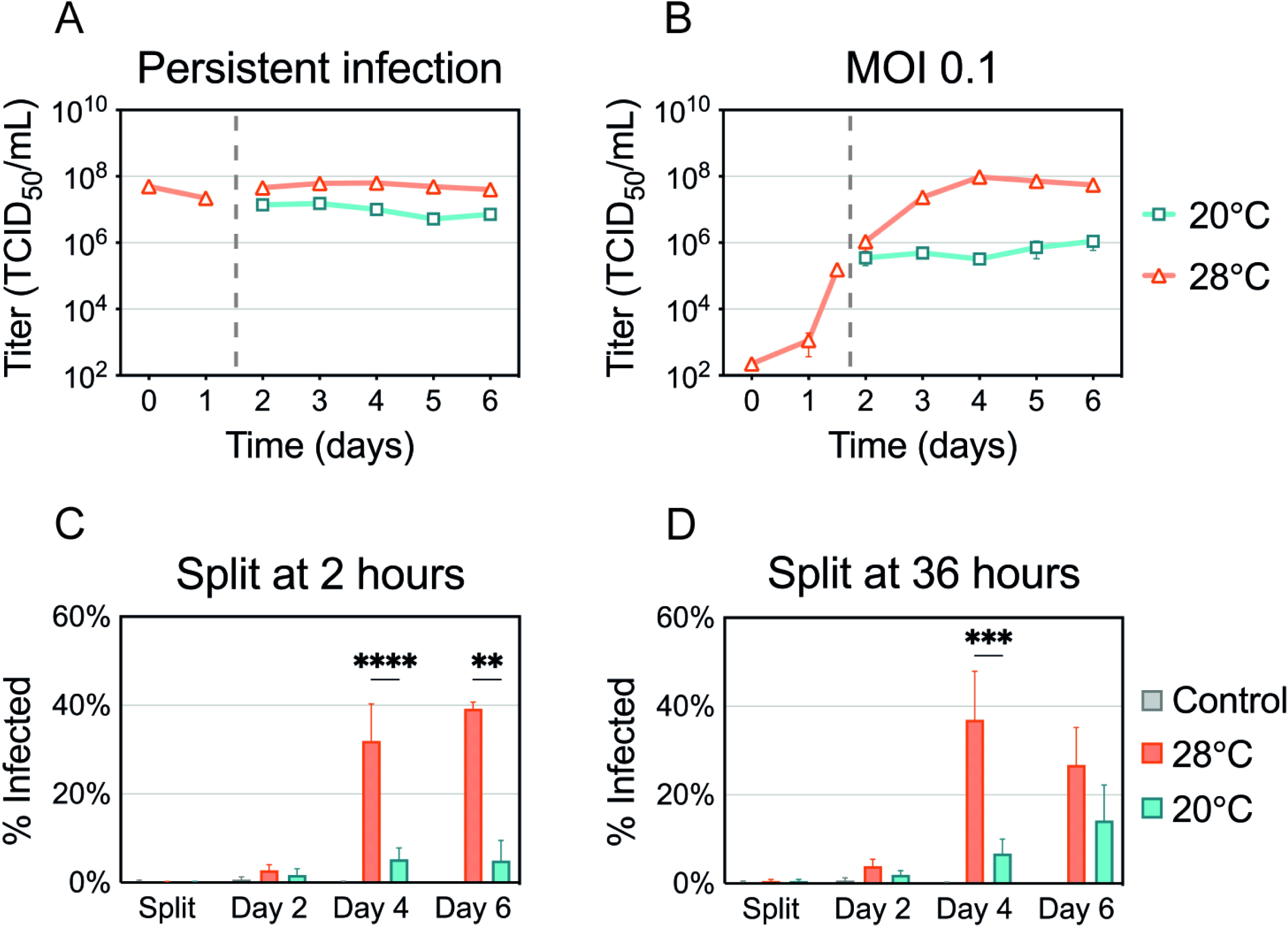
ZIKV production and spread at suboptimal temperature. (**A**) C6/36 cells with established ZIKV infection were split and incubated at 20°C or 28°C for 5 d. (**B**) C6/36 cell infected at an MOI of 0.1 were split at 36 h p.i. and incubated at 20°C and 28°C for 5 d. Supernatants were collected and replaced daily, and virus titers were determined. Data shown for persistent ZIKV infection are means ± SEM of results from two biological replicates each consisting of three technical replicates (n=6). Data shown for MOI 0.1 are means ± SEM of results from three independent experiments. Dashed lines represent when the cells were split. (**C and D**) Percentage of ZIKV-positive cells was assessed using flow cytometry. (**C**) Cells were infected with ZIKV (MOI 0.1) for 2 h at 28°C and then split and incubated at 20°C or 28°C. (**D**) Cells infected with ZIKV were kept at 28°C for 36 h and then split and incubated at 20°C or 28°C. ZIKV-positive cells were measured after splitting (2 or 36 h p.i.) and 2, 4, and 6 d p.i. Data shown are means ± SEM of results from seven independent experiments. Statistical differences were assessed by a two-way ANOVA with Bonferroni’s correction where cells at 20°C were compared to cells at 28°C at every time point. ** *p* < 0.01, *** *p* < 0.001, **** *p* < 0.0001

Next, we infected C6/36 cells with a low MOI (0.1) and incubated them at 28°C for 36 hours, during which ZIKV infection was established in a small proportion of cells (Fig 4B). The cells were then split and incubated at 20°C or 28°C for five additional days. While virus titers increased up to three logs at 28°C, the titers plateaued in C6/36 cells maintained at 20°C. This plateau could be the result of inhibited viral spread from the previously infected cells to naïve cells, or an overall reduction in virus production in both cells with established infection and newly infected cells.

To determine if the plateau in virus titers was due to inhibition of virus spread, we quantified the number of infected cells over time. We infected cells with a low MOI and incubated them at 28°C for either two hours, to mimic the original replication curves (Fig 4C), or 36 hours, to mimic the later experiment (Fig 4D). The cells were then maintained at 20°C or 28°C, stained for ZIKV antigen, and analyzed using flow cytometry. We saw that cultures maintained at 20°C displayed low levels of ZIKV-positive cells, suggesting the virus was unable to spread and replicate in naïve cells (Fig 4C). When the cells were maintained at 28°C for 36 hours before shifting, more ZIKV-positive cells were detected at 20°C; however, 28°C enhanced the production of ZIKV antigens. Taken together, these data suggest that the cells with an established infection are capable of producing infectious particles, however the virus produced is not able to efficiently spread and establish infection in uninfected cells at 20°C.

### ZIKV entry kinetics

In order to elucidate the kinetics of ZIKV internalization and fusion in C6/36 cells, infected cells were treated with chlorpromazine and ammonium chloride. Chlorpromazine (CPZ) inhibits clathrin-mediated endocytosis, an entry pathway used by ZIKV in mammalian cells (26, 27). Ammonium chloride (NH_4_Cl) is a weak base that increases the endosomal pH and prevents the low pH-dependent conformational changes in the E protein required for fusion (28–30). To ensure these compounds inhibit ZIKV entry in C6/36 cells, we first confirmed both drugs reduced viral titers while remaining non-cytotoxic (Fig 5A and 5B). We then performed a time-of-addition assay with NH_4_Cl to establish the dynamics of internalization and fusion steps in C6/36 cells (Fig 5C). We treated the cells with NH_4_Cl at the indicated time points to determine when ZIKV completes fusion and entry in cells maintained at 28°C. If NH_4_Cl was added one hour after infection, it blocked 65% of the virus produced in the mock control, while only 25% of the virus was inhibited if added at two hours. This suggests that most ZIKV particles enter C6/36 cells within the first two hours of infection at 28°C.

**FIG 5.**
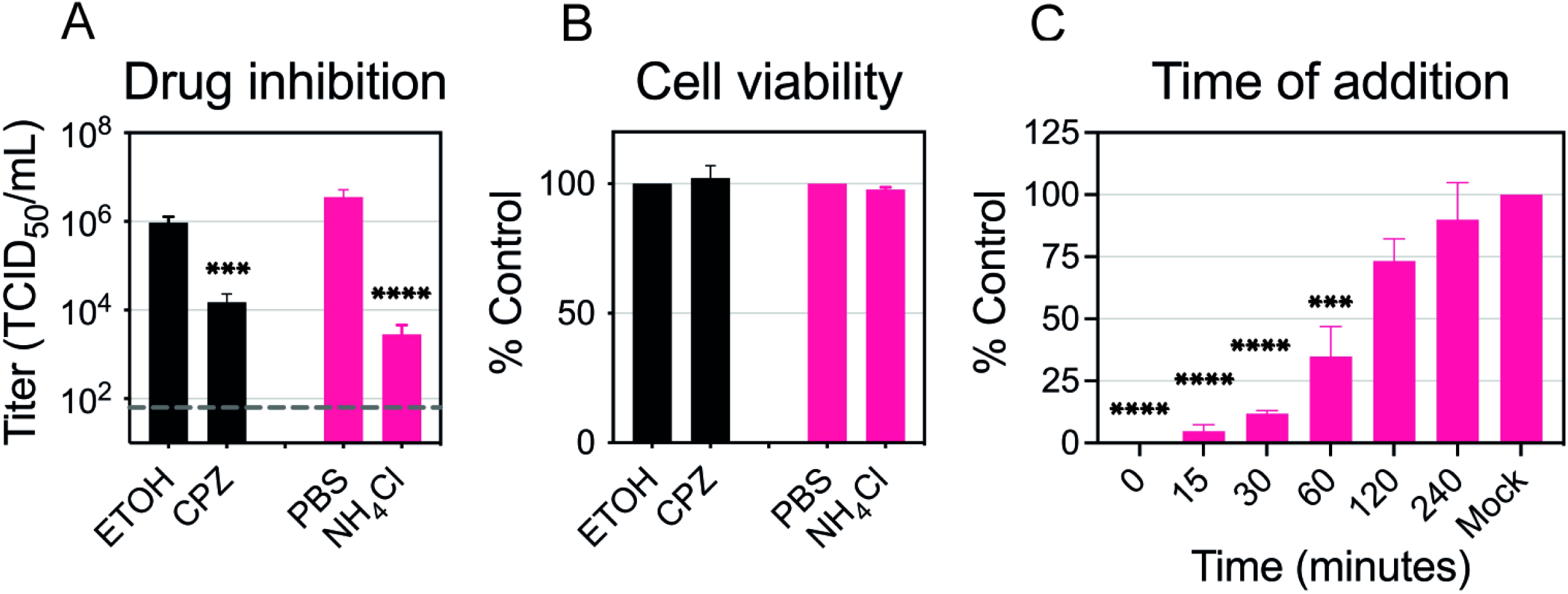
ZIKV entry dynamics in C6/36 cells. (**A**) Drug inhibition assay. CPZ (50 μM) and NH_4_Cl (25 mM) were added 2 h prior to ZIKV infection (MOI 0.1) and were kept during infection. After a 2 h incubation at 28°C, the infectious media with the drug or vehicle was removed and replaced with fresh media. Supernatants were collected at 48 h p.i. and titrated. Dashed line represents the limit of detection. (**B**) Cell viability. The cells were treated with the same concentrations of the drug or the same amount of vehicle for 4 h and ATP levels were measured after 2 days. Data are presented as mean ± SEM from three independent experiments for each drug. Statistical differences were assessed by using a two-way ANOVA followed by Bonferroni’s test for multiple comparisons. (**C**) Time-of-addition. NH_4_Cl (25 mM) was added to the ZIKV-infected cells at the following time points: 0-, 15-, 30-, 60-, 120-, and 240-min p.i. and was kept for 4 h. Supernatants were collected at 48 h p.i. and titrated. Cell titers were normalized to mock infection and data are presented as mean ± SEM from three independent experiments. Statistical significance was determined by using a one-way ANOVA on log transformed data with Dunnett’s correction where mock was compared to the time of drug-addition. * *p* < 0.05

### Effects of suboptimal temperature on ZIKV entry

Next, we wanted to determine when the cool temperature is most affecting ZIKV replication. We evaluated the effect of transferring infected cells between suboptimal (20°C) and optimal (28°C) environments on viral titers. If the cells were infected and maintained at 20°C for two or six hours before being shifted to 28°C, the virus titers were similar to cells maintained at 28°C for the entire time period (Fig 6A). Based on the previously established entry dynamics, this suggests that ZIKV entry was not inhibited at the suboptimal temperature. However, if cells were infected at 20°C and maintained at the cool temperature for 12 hours before shifting to 28°C, fewer particles were produced. When infected cells were shifted from warm (28°C) to cool (20°C) environments, we found that considerably fewer particles were produced if the shift occurred at any point within the first 12 hours after infection (Fig 6B). This suggests that ZIKV requires more than 12 hours at 28°C to establish a productive infection. In combination with our previous findings (Fig 4 and Fig 5), these data suggest that cool temperatures block efficient replication after fusion, but late stages of the virus replication cycle in cells with established infection can proceed at cool temperatures.

**FIG 6.**
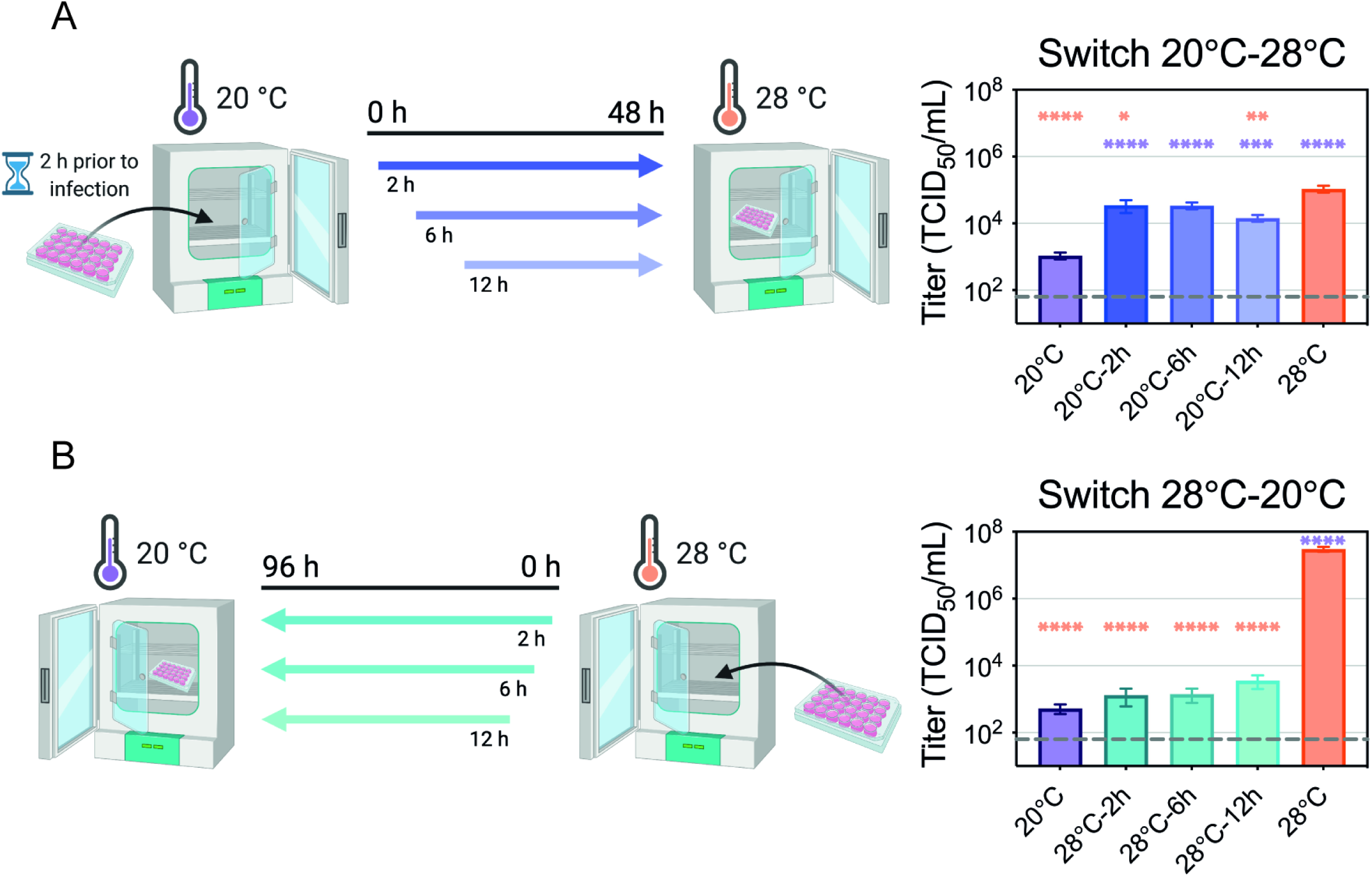
ZIKV entry at suboptimal temperature. (**A**) Cool-to-warm temperature switch. C6/36 cells (6 × 10^5^ cells/mL) were plated and incubated at 20°C 2 h prior to infection. Cells were then inoculated with ZIKV (MOI 0.1) for 2 h and moved to 28°C at 2-, 6-, or 12- h p.i. As a control, one set of cells was maintained at 20°C while another was maintained at 28°C throughout the experiment. Supernatants were collected at 48 h p.i. and titrated. (**B**) Warm-to-cool temperature switch. C6/36 cells were plated at a same density and infected with ZIKV (MOI 0.1) for 2 h. The cells were transferred from 28°C to 20°C at 2-, 6-, or 12- h p. i. As a control, one set of cells was maintained at 28°C while another was maintained at 20°C throughout the experiment. Supernatants were collected at 96 h p.i. and titrated. Dashed lines represent the limit of detection. Experimental designs were created with BioRender.com. Data are presented as mean ± SEM from three replicates for each experiment. Statistical differences were determined by a one-way ANOVA on log transformed data followed by Dunnett’s test for multiple comparisons where each condition was compared to both 20°C (purple significance symbol) and 28°C (orange significance symbol) controls. * *p* < 0.05, ** *p* < 0.01, *** *p* < 0.001, **** *p* < 0.0001

### Effects of suboptimal temperature on virus recovery, translation and dsRNA production

Our data indicate that cooler temperature primarily inhibits ZIKV replication after the initial two-hour viral entry period. To further confirm the phenotype, we bypassed binding, internalization, fusion, and nucleocapsid disassembly steps by transfecting full-length genomes into the cells and quantifying the amount of infectious viral particles produced (Fig 7A). If the cells were transfected and maintained at 28°C, infectious ZIKV particles were readily detected after two days. However, if the cells were transfected at 28°C and transferred to 20°C one hour following transfection, no infectious virus was produced. If 20°C-adapted cells were transfected and kept at 20°C no virus was produced, but if transferred to 28°C one hour following transfection, virus was produced at similar levels to the cells maintained at 28°C.

**FIG 7.**
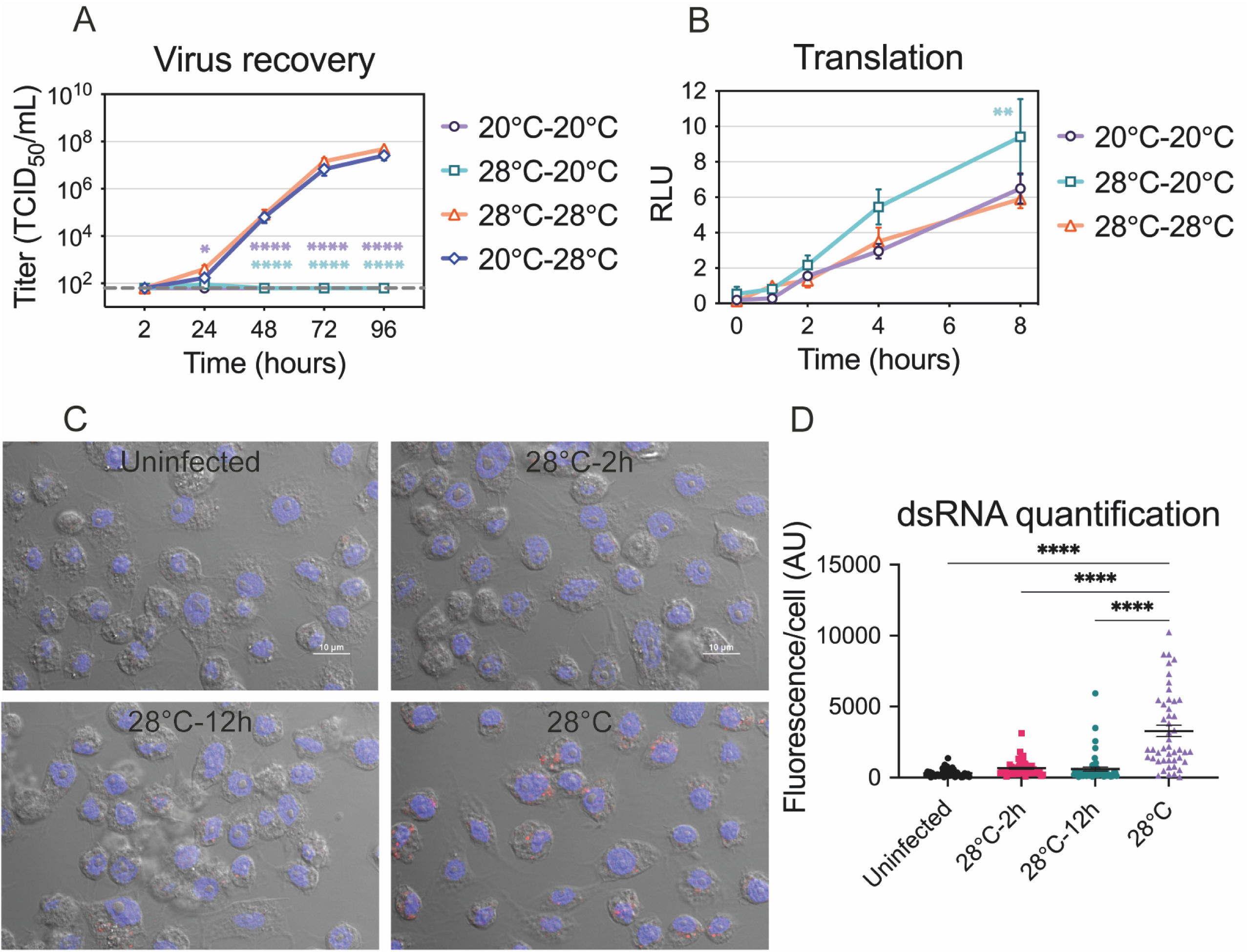
Virus recovery, translation, and dsRNA production at suboptimal temperature. (**A**) Virus recovery. Standard and 20°C-adapted cells were transfected with infectious ZIKV RNA at their respective temperatures for 1 h. After transfection, one plate with standard cells was kept at 28°C and one was transferred to 20°C, and one plate with 20°C-adapted cells was kept at 20°C and one was transferred to 28°C. At 2-, 24-, 48-, 72- and 96-h after transfection, supernatants were collected and titrated. Dashed line represents the limit of detection. Data are presented as mean ± SEM from at least three independent experiments. Statistical significance was determined by using a two-way ANOVA on log transformed data followed by Dunnett’s multiple comparisons test. For each time point, standard cells kept at 28°C (28°C-28°C) were compared to standard cells kept at 20°C (28°C-20°C), 20°C-adapted cells kept at 20°C (20°C-20°C), and 20°C-adapted cells kept at 28°C (20°C-28°C). (**B**) Translation. Standard and 20°C-adapted C6/36 cells were transfected in duplicates with a translation reporter construct at their respective temperatures. One plate with standard cells was kept at 28°C and one was switched to 20°C 1 h after transfection, while 20°C-adapted cells were kept at 20°C. At 0-, 1-, 2-, 4-, and 8-h after transfection, the cells were lysed and stored until all time points were collected. Luminescence was normalized to 28°C control at 1 h after transfection. Data shown are means ± SEM from three independent experiments each performed in duplicate wells. Statistical differences were assessed by a two-way ANOVA using Dunnett’s test for multiple comparisons. For each time point, standard cells kept at 28°C (28°C-28°C) were compared to standard cells kept at 20°C (28°C-20°C) and 20°C-adapted cells kept at 20°C (20°C-20°C). (**C**) Accumulation of dsRNA intermediates. C6/36 cells were infected with a high MOI of ZIKV (MOI >10) at 28°C and were shifted to 20°C at 2- or 12- h p. i. Cells were processed for immunofluorescence using a monoclonal antibody to detect dsRNA at 48 h p. i. Uninfected cells were used as a control for specificity of the antibody against dsRNA. Nuclei are shown in blue and dsRNA intermediates are shown in red. (**D**) Quantification of dsRNA fluorescence signal in ZIKV-infected cells as described in panel C. Points represent individual cells analyzed (n = 47 to 53). Statistical significance was determined by using a one-way ANOVA with Dunnett’s correction where cells infected and kept at 28°C were compared to each condition. AU, arbitrary units. The color of the significance symbol corresponds to the color for each condition. * *p* < 0.05, ** *p* < 0.01, **** *p* < 0.0001

Efficient translation is required for virus to be produced from the transfected genomes. To directly measure translation efficiency, we designed a translation reporter construct containing a luciferase coding region flanked by the 5’ and 3’ ZIKV untranslated regions and transfected the RNA into the cells (Fig 7B). There was no difference in luciferase signal between the cells that were transfected and maintained at 28°C and the 20°C-adapted cells that were transfected and maintained at 20°C. Interestingly, there was a slight increase in luciferase production in the cells that were transfected at 28°C and incubated at 20°C for eight hours.

Since translation was not inhibited by the cool temperature, we wanted to investigate the effect of the cool temperature on genome replication (Fig 7C). Genome replication was detected by staining cells with an antibody that detects double-stranded RNA (dsRNA), an intermediate produced during genome replication. When ZIKV-infected cells were transferred to 20°C two or 12 hours after transfection, we were unable to detect dsRNA intermediates at the cool temperature after 48 hours. Taken together, our data suggest that genome replication is inefficient at cool temperatures in mosquito cells.

### Replication curves of different ZIKV strains at suboptimal temperature

Lastly, we wanted to explore if limited virus replication in C6/36 cells at 20°C was common among different ZIKV isolates or lineages. We compared replication curves of ZIKV Mex 1-44 to another Asian-lineage, SPH (Fig 8A), and to two African-lineages, MR766 and IbH (Fig 8B and 8C). As previously observed with Mex 1-44 (Fig 1A), SPH replication was reduced at 20°C; low levels of virus were detected on day six and only slightly increased through day 10. In contrast, both African-lineage strains replicated faster, with detectable replication occurring earlier at day four and viral production at both 20°C and 28°C reaching similar values over the course of the experiment.

**FIG 8.**
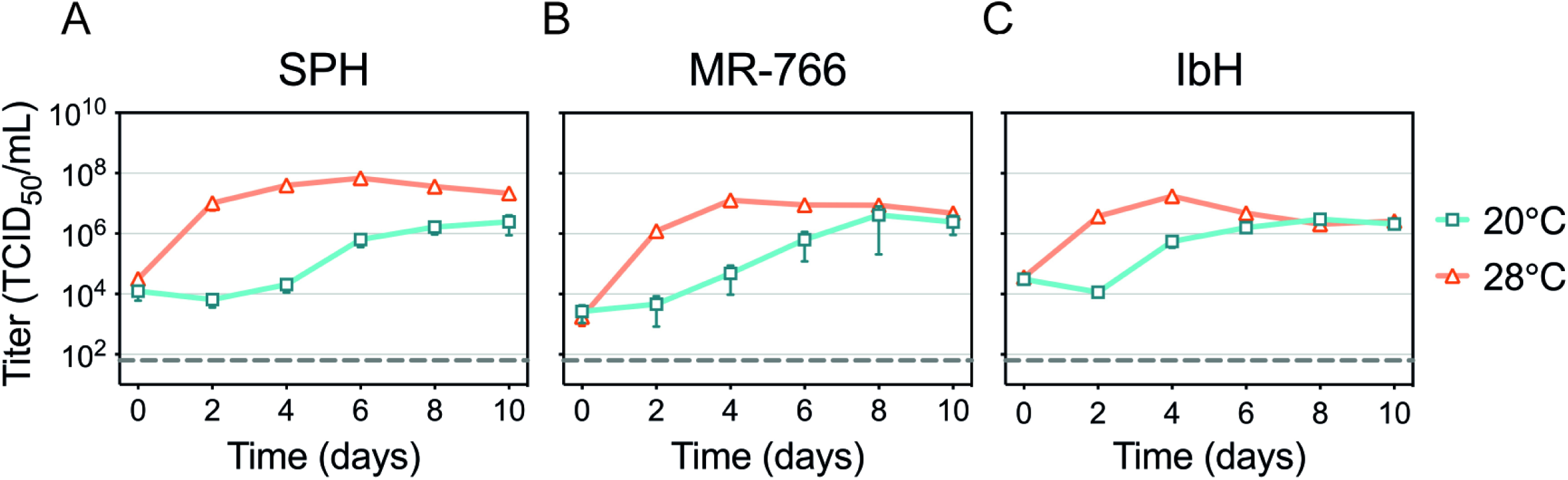
Replication of different ZIKV strains at sub-optimal temperatures. Replication rates for (**A**) SPH, (**B**) MR766, and (**C**) IbH at cool temperature. The cells were infected at an MOI of 0.1 for 2 h at 28°C and incubated at either 20°C or 28°C for 10 d. Supernatants were collected every 48 h and titrated. Dashed lines represent the limit of detection. Data shown are means ± SEM of results from five replicates for each virus.

## Discussion

In order to be successfully transmitted, arboviruses need to adapt to distinct hosts and persist within a wide temperature range. Environmental temperature plays a critical role in arboviral transmission since mosquito vectors are susceptible to changes in temperature (9, 31–35). We have previously demonstrated that ZIKV transmission in *Ae. aegypti* mosquitoes is optimized at 29°C with a thermal range of 22.7°C-34.7°C (25). Low ambient temperatures, that are well within the range for survival of *Aedes* species, are known to inhibit arboviral replication and transmission (12, 36). However, the molecular mechanisms underlying the effects of temperature on virus replication efficiency and vector competence remain unclear. Here, we present data that focuses on how temperature alters ZIKV replication *in vitro* in C6/36 cells to characterize this relationship in a minimal system that excludes systemic host responses. We hypothesized that suboptimal temperatures alter the intracellular environment, which consequently inhibits virus replication, and/or that temperature directly inhibits virus replication. First, we showed that ZIKV replication is inhibited at suboptimal temperatures even when cells are adapted to those temperatures and therefore not undergoing acute temperature stress. We further clarified the steps of the virus replication cycle that are most affected by cool temperatures, examining viral entry, translation, and genome replication. We found that cool temperatures did not inhibit ZIKV entry or translation, but it significantly reduced the production of dsRNA replication intermediates (Fig 9A and B). However, if cells with an established infection were shifted to 20°C, all the subsequent steps were efficiently completed, suggesting that cool temperatures decrease the efficiency of genome replication (Fig 9C).

**FIG 9.**
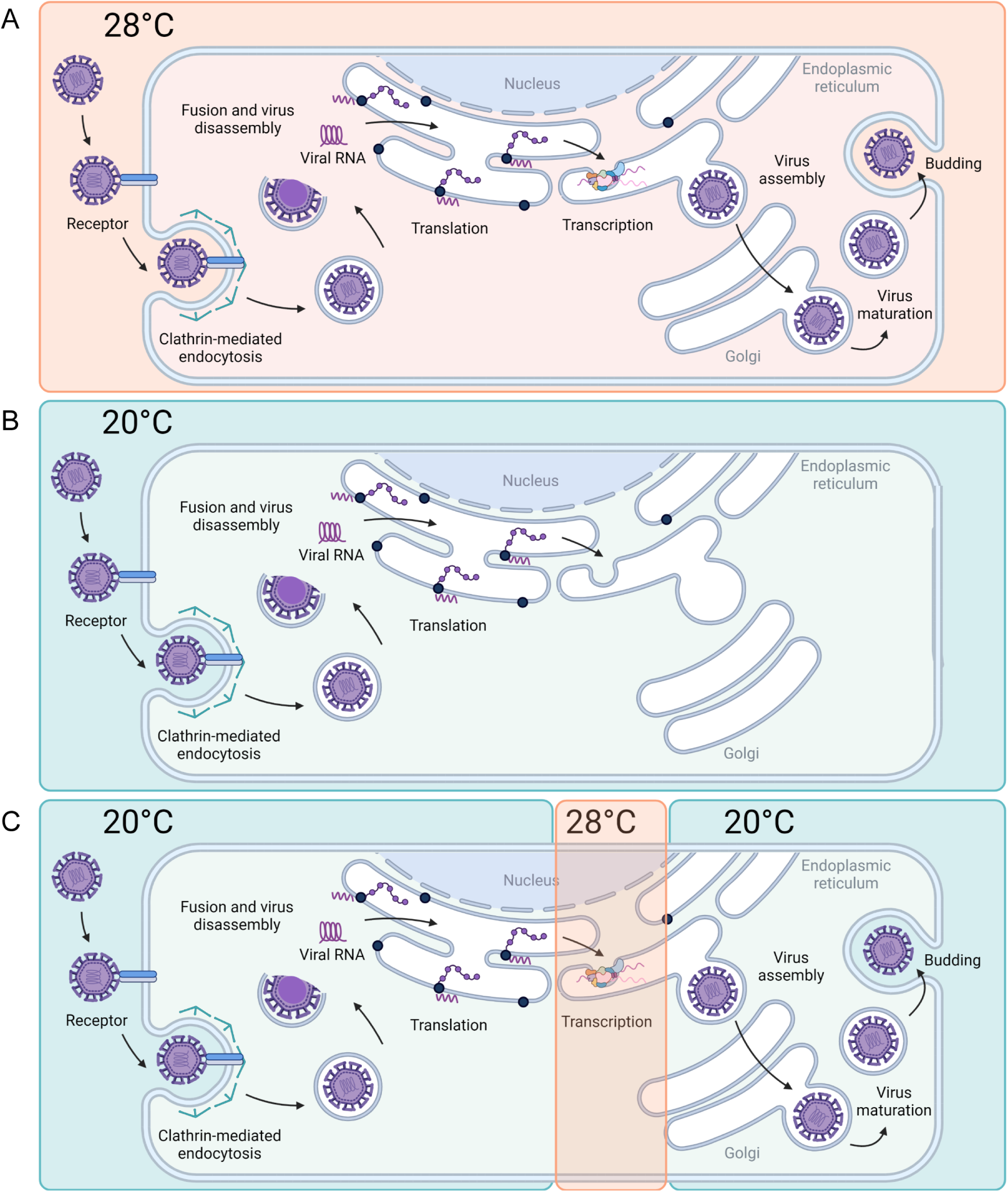
Model of ZIKV replication cycle at 20°C and 28°C. Replication of ZIKV efficiently occurs at 28°C, completing all steps (A). When ZIKV infection is initiated at 20°C, infection is stalled after capsid disassembly, but before dsRNA intermediates can be detected (B). Late stages in the ZIKV replication cycle can occur at 20°C if RNA replication steps occur at 28°C (C).

Temperature is known to induce molecular changes that impact structures and functions of proteins, lipids and nucleic acids, which in turn can affect intracellular processes, enzymatic activity, membrane fluctuations, cell metabolism, viability, and induce temperature-shock responses. Therefore, temperature modifies virions and their interactions with cellular components, and is one of the most important factors impacting infection dynamics during viral replication (37). Although arboviruses can tolerate drastic temperature changes, replicating efficiently in both mammalian (∼37°C) and mosquito (∼28°C) hosts, we found that exposing ZIKV-infected mosquito cells to a range of temperatures affected virus yields over time. During early infection, we observed slower replication kinetics at lower temperatures and faster replication at higher temperatures (Fig 1E). These faster replication rates of ZIKV at high temperatures could be caused by increased cellular metabolism, enzymatic activity, and fluidity of cellular membranes. Moreover, there is evidence that heat-shock response can facilitate entry and replication of Zika, dengue, West Nile, and Japanese encephalitis viruses (38–40). It has also been suggested that high temperatures have direct influence on viral RNA structure and long-range RNA-RNA interactions (21, 41) as well as on the conformation of RNA-dependent RNA-polymerase (20) resulting in enhanced replication and higher titers. Later in infection, we observed decreased viral titers at cool and high temperatures, and increased titers around intermediate temperatures (Fig 1F). The drop in virus yields observed at high-temperature conditions was caused by a prolonged heat-shock response that affected cell viability. Cool temperatures, on the other hand, resulted in low virus yields at all time points. Although cold temperatures decrease cellular metabolism, enzymatic activity, and membrane fluidity (42, 43), none of these potential alterations in the intracellular environment inhibited DENV or CHIKV replication in C6/36 cells at 20°C to the same extent that ZIKV replication was inhibited, suggesting ZIKV is more sensitive to temperature than the other viruses.

First, we evaluated ZIKV entry, which includes virus-cell attachment, internalization, and envelope-cell membrane fusion, at 20°C. ZIKV enters the cells through clathrin-mediated endocytosis and requires low pH for fusion (26). By adding ammonium chloride at different times after ZIKV infection, we established that most ZIKV virions enter C6/36 cells within two hours at 28°C. Since cells were infected at 28°C for two hours before shifting to other temperatures in the majority of the experiments, we suggest the temperature inhibition was preventing a step in the replication cycle following membrane fusion. Although temperature is known to affect viral structure (16, 17, 44), the stability of virus particles (45, 46), and alter viral protein conformational changes necessary for virus entry (18, 19, 24), incubating the cells at 20°C during the first six hours of infection did not affect virus yields (Fig 6A), suggesting the cool temperature is not blocking early steps in the replication cycle (Fig 9). Moreover, transfecting the cells with viral RNA, thereby bypassing entry and nucleocapsid disassembly steps, resulted in an inhibition of virus production similar to infecting the cells at 20°C (Fig. 7A). However, while transfecting viral RNA into cells maintained at 20°C prevented virus production, transfecting a translation reporter construct demonstrated that translation is not affected by cool temperature (Fig. 7B). When we specifically investigated genome replication, we could not readily observe dsRNA replication intermediates at cool temperature. Late steps in virus replication cycle, such as budding, maturation, and viral egress, are not likely disrupted by cool temperature since the cells with an established infection can produce similar number of infectious particles at 20°C and 28°C (Fig. 4A).

There are several steps viruses undergo after entry and before genome replication that could be sensitive to cooler temperature. After translation and polyprotein processing, ZIKV must induce extensive ultrastructural changes and rearrangement of the intracellular membranes to provide the platforms required for the formation of replication complexes (47, 48). Replication inside membrane compartments concentrates the molecules necessary for replication and particle assembly, but also helps evade host antiviral defense mechanisms (49). The formation of cytoplasmic membranous vesicles has been well-demonstrated among positive-sense RNA viruses including *Picornaviridae*, *Togaviridae*, and *Flaviviridae* families. Several picornaviral nonstructural proteins (2C, 2BC, and 3A) are known to be required for formation of membranous vesicles (50, 51). The endoplasmic reticulum (ER) is the primary site for flavivirus replication, and DENV is known to perturb lipid homeostasis in mosquito cells (52) and to reprogram the ER protein synthesis and processing environment to promote viral replication (53). Studies have shown that DENV utilizes fatty acid biosynthesis resulting in lipid enrichment that has the capacity to destabilize and change the curvature and permeabilization of membranes (52) and NS3 protein was identified to redistribute fatty acid synthase (FAS) to the site of viral replication (54). Further, a drug associated with the modulation of the host cell lipid metabolism (PF-06409577) inhibited ZIKV, DENV and WNV infection (55). It is well known that cool temperatures can affect membrane fluidity, but it remains unclear how temperature could alter the ability of viral non-structural proteins to interact with cellular factors and disturb membrane rearrangement.

RNA cyclization, another posttranslational step in virus replication cycle, is known to be temperature dependent and it is a prerequisite for the initiation of flavivirus replication. Highly structured 5’- and 3’-untranslated regions (UTRs) support a switch in the RNA genome structure from a linear to a cyclized form which enables NS5 polymerase to initiate RNA replication. Studies have shown that cultivation temperature decisively determines the replication efficiency of the WNV and ZIKV RNA in mammalian and mosquito cells (21, 41). WNV long-range RNA-RNA interactions *in vitro* are enhanced at high temperatures (37°C), while at low temperatures (28°C) genome cyclization is less efficient and therefore replication starts at later time points. It has been suggested that this “RNA thermometer” activity could be involved in modulating the replication efficiency during host switching (21). A study by X. D. Li et al. has shown that a single-nucleotide mutation in the DAR (downstream of AUG region) is sufficient to abolish long-range RNA-RNA interactions and *de novo* RNA synthesis of ZIKV at 28°C but not at 37°C. A more restricted DAR complementarity for viral replication at lower temperatures is additional evidence that temperature is a critical determinant of replication kinetics. Authors suggested that a temperature-specific function of the DAR element enables ZIKV to resist the temperature stresses of different hosts (41). Another study suggested that the active site of the DENV RNA-dependent RNA-polymerase (RdRp) exists in two conformational states, a more rigid conformation at lower temperatures and a more mobile conformation at higher temperature, and that temperature determines how the enzyme recognizes the conformation of the template RNA and affects RNA synthesis (20).

Interestingly, in this study, low temperature affected Asian-lineage ZIKV isolates (Mex 1-44 and SPH) and African-lineage ZIKV isolates (MR766 and IbH) differently. While Mex 1-44 and SPH replicated at a slower rate and had lower virus titers at 20°C, MR766 and IbH replicated faster, and titers peaked at similar levels at 20°C and 28°C. African and Asian lineages were previously shown to have different phenotypes in cell culture, *Ae. aegypti* mosquitoes, and chicken embryos (56). While African isolates have faster replication rates in C6/36 cells and induce higher mortality in embryos than Asian-lineage isolates, they seem to be less infectious to *Ae. aegypti* mosquitoes (56). Moreover, ZIKV isolates from Asian and African lineages displayed different replication kinetics, cytopathic effects, and impacts on human neural progenitor cell function (57, 58), as well as the ability to infect human placental trophoblast (59), cause severe brain damage (60), *in utero* infection in pigs (61), and disease progression in mice (62). Comparative analysis of viral entry showed that divergent ZIKV strains enter cells in a highly conserved manner (30), suggesting genetic variation between ZIKV isolates affects some other aspect of ZIKV replication cycle with consequences on infection kinetics and pathogenesis.

We have shown that temperature affects flavivirus and alphavirus replication kinetics differently. ZIKV and DENV responded similarly to temperatures; however, the extent to which temperature affected virus yields differed. DENV replication was more efficient and reached higher titers at cool temperatures than ZIKV, which is consistent with our *in vivo* study showing the predicted thermal minimum for DENV is 5°C lower than that of ZIKV in *Ae. aegypti* mosquitoes (25) and even lower in *Ae. albopictus* mosquitoes (11). While flavivirus titers increased with the increase in temperature early during infection, CHIKV titers were the highest at intermediate temperatures and lower at both low and high temperatures. Although attenuated CHIKV strain replicated significantly better at cool temperatures and eventually reached almost same peak titers at 16°C and 28°C *in vitro*, studies *in vivo* showed that the low ambient temperature of 18°C may impede transmission of some CHIKV strains by *Ae. albopictus* (63). The same group further characterized gene expression in mosquitoes during CHIKV infection at 18°C, 28°C, and 32°C and reported that FAS was among the topmost downregulated gene at low temperature (64). As discussed above, FAS is important for the formation of replication complexes and its downregulation could affect virus replication and transmission. However, another study showed that high viral titers are not sufficient for virions to cross the midgut escape barrier. It suggested that viral replication kinetics are more important for midgut escape and transmission ability of the CHIKV (65), since slowly replicating variants are less able to overcome the midgut escape barrier. This could explain the block in transmission of CHIKV at 18°C (63) and ZIKV at 20°C (25).

After exposing *Ae. albopictus*-derived C6/36 cells infected with ZIKV to the same temperatures we previously exposed ZIKV-infected *Ae. aegypti* mosquitoes (25), we observed similar qualitative trends. High temperatures caused faster replication rate *in vitro*, as well as higher proportion of infected mosquitoes and shorter EIP *in vivo*, while affecting cell viability and mosquito survival. Low temperatures caused a significant reduction in viral titers *in vitro* and a low proportion of infected mosquitoes, longer EIP, and impaired transmission *in vivo*. We have also investigated the transcriptome profile of ZIKV-infected mosquitoes at low and high temperatures and showed that temperature dramatically shapes mosquito gene expression (66). Genes that were mostly altered at cool temperature involved aspects of blood-meal digestion, ROS metabolism, and mosquito innate immunity. Although temperature causes systemic changes in mosquitoes that can alter virus replication, the direct effect of temperature on ZIKV replication was still visible at the cellular level. While the effects of temperature on ZIKV transmission and vector competence have been well documented (7, 25, 67–69), R. A. Murrieta et al. recently demonstrated that temperature also impacts virus evolution. They showed that temperature significantly modifies selective environment within mosquitoes, and that fluctuating temperatures cause strong purifying selection mostly in ZIKV NS5 (70). Future experiments should thus characterize the phenotypes at suboptimal temperatures and investigate potential evolutionary patterns associated with ZIKV replication at cool temperatures.

Understanding transmission dynamics and epidemiology of infectious diseases is crucial for successfully controlling current and preventing future outbreaks. Characterizing the link between pathogen replication and environmental conditions is important to better understand how environmental factors shape transmission.

## Materials and Methods

### Cell lines

C6/36 mosquito cells (ATCC CRL-1660) from *Ae. albopictus* were maintained in Leibovitz’s L-15 medium supplemented with 10% fetal bovine serum (FBS) at 28°C. Some C6/36 cells were adapted to grow at 20°C. To adapt the cells, they were initially maintained in L-15 medium with the higher FBS content (20%). After the cells were maintained at 20°C for several weeks, the amount of FBS was gradually reduced to 10% over several months. Vero (African green monkey kidney) cells were cultured in Dulbecco’s Modified Eagle Medium (DMEM) with 5% FBS at 37°C, 5% CO_2_.

### Virus isolates and viral titer determination

All viral stocks were generated in Vero cells and tested negative for *Mycoplasma* contamination using a MycoSensor PCR Assay Kit (Agilent). Detailed passage histories of the ZIKV isolates used in this study were previously described (56). ZIKV Mex 1-44 was isolated from a field-caught *Ae. aegypti* mosquito in Chiapas, Mexico, in 2016, and obtained from the University of Texas Medical Branch (71). ZIKV SPH originated from a Brazilian clinical sample in 2015 and was obtained from the Oswaldo Cruz Foundation (72). Two African-lineage isolates, ZIKV MR766 and IbH, were purchased from American Type Culture Collection (catalog number VR-1838™ and VR-1829™, respectively) (73). The DENV-2 isolate (PRS 225488) originated from human serum collected in Thailand in 1974 and was obtained from the World Reference Center for Emerging Viruses and Arboviruses at the University of Texas Medical Branch. Attenuated CHIKV strain (181/c25) was generated as previously described (74). Viral RNA was produced by *in vitro* transcribing pSinRep5-181/25ic, a gift from Terence Dermody (Addgene plasmid #60078) (75). Capped RNA was transfected into 293T cells and virus containing supernatants were collected when cells showed signs of cytopathic effect. All viral titers were determined by endpoint dilutions on Vero cells. The tissue culture infectious dose 50 (TCID_50_) was determined using the Spearman-Karber method (76).

### Virus replication curves

C6/36 cells were seeded in 12-well plates at a density of 4 × 10^5^ cells/mL several hours prior to infection. Cells were inoculated with ZIKV using a multiplicity of infection (MOI) of 0.1, DENV with an MOI of 0.1, or CHIKV with an MOI of 0.005 for two hours. The cells and media were kept at the indicated constant temperatures for 10 days. Every 24 or 48 hours, half of the supernatant was collected and replaced with fresh media equilibrated to each temperature. Similarly, Vero cells were plated at a density of 2.5 × 10^5^ cells/mL and inoculated with ZIKV (MOI 0.1) for one hour at 37°C. The cells were incubated at different constant temperatures, and half of the cell supernatant was collected every 24 hours for seven days.

### Cell proliferation

C6/36 cells were stained using the CellTrace™ Violet Cell Proliferation Kit (Invitrogen) as per manufacturer’s instructions and plated at a density of 4 × 10^5^ cells/mL in 24-well plates. The cells were maintained at different constant temperatures and intracellular fluorescence, which decreases with every generation, was determined using flow cytometry at indicated time points. The mean fluorescence intensities (MFI) were normalized to the MFI value determined at the time of staining to calculate the number of divisions that occurred.

### Cell viability

Cell viability was monitored using the CellTiter-Glo® Luminescent Cell Viability Assay (Promega) and a Glomax® Explorer (Promega) plate reader as per the manufacturer’s instructions. C6/36 cells were plated at a density of 4 × 10^5^ cells/mL in 24-well plates and treated with the virus, temperature, or drug. At indicated time points, cell viability was determined and compared to indicated controls.

### pAc5-STABLE2-neo transfection

C6/36 cells were seeded in two T-25 flasks at a density of 6 × 10^6^ cells per flask the night before transfection. One flask was transfected with Ac5-STABLE2-neo plasmid, a gift from Rosa Barrio and James Sutherland (Addgene plasmid #32426) (77), using Lipofectamine™ 3000 Transfection Reagent (Invitrogen) as per manufacturer’s instructions. The plasmid encodes for Flag-mCherry, GFP, and Neo^R^ under the control of the insect Actin5C promotor. Six hours after transfection, both transfected and control cells were lifted, plated in six 24-well plates, and transferred to the indicated constant temperature incubators. Cells were resuspended in PBS and GFP production was measured using flow cytometer (BD-LSRII) 24 and 72 hours after temperature exposure.

### ZIKV production during temperature shift experiments

Persistently infected ZIKV C6/36 cells were produced by maintaining an infected culture in a T-75 flask at 28°C for several weeks. To monitor viral production, 1 mL of cell supernatant was collected on two consecutive days. The cells were then seeded in two 6-well plates at a density of 6 × 10^5^ cells/mL. One plate was maintained at 28°C and the second was placed at 20°C. Every 24 hours, the entire cell supernatant was collected and replaced to measure the daily virus production.

For the experiment with actively spreading infection, C6/36 cells were seeded in a T-25 flask at a density of 6 × 10^5^ cells/mL. The following day, the cells were infected with ZIKV (MOI 0.1) for two hours at 28°C. After incubation, infectious media was removed, and the cells were washed once before fresh media was added. Supernatant was collected at the indicated time points. After the 36-hour time point sample was collected, the cells were washed and seeded in two 6-well plates. Each plate was incubated at either 20°C or 28°C for five days. Every 24 hours, the entire cell supernatant was collected and replaced to measure the daily virus production.

### Detection of ZIKV-positive cells at optimal and suboptimal temperatures using flow cytometry

C6/36 cells were plated in three T-25 flasks at a density of 2 ×10^6^ cells/mL. Two flasks were infected with ZIKV (MOI 0.1) for two hours at 28°C. Cells from one flask were split into two 6-well plates and transferred to 20°C or 28°C at 2 hours following infection. Infected and uninfected control cells from the other two flasks were kept at 28°C and were split and transferred 36 hours following infection. The cells were stained for intracellular ZIKV protein NS1 and the percentage of ZIKV-positive cells was determined using flow cytometer at the indicated time points. Briefly, the cells were resuspended in fresh media, fixed with 4% paraformaldehyde for 15 minutes and permeabilized with 0.1% Triton X-100 in 1×PBS for 15 minutes. The cells were centrifuged at 300 × g for 10 minutes and resuspended in the primary antibody (anti-ZIKV NS1 mouse MAb, BioFrontTech) diluted in permeabilization buffer at a concentration of 1:500. After 45 minutes of incubation at 37°C, the cells were washed three times in permeabilization buffer by centrifuging for 5 minutes at 300 × g. After the last spin, the cells were resuspended in the secondary antibody (Alexa Fluor™ 647-conjugated goat anti-mouse Ab, Invitrogen) at a concentration of 1:1000, and incubated for 30 minutes at 37°C. The cells were washed three times and resuspended in 1×PBS. The percentage of infected cells for each treatment was determined for 20,000 single cells.

### Drug inhibition and time-of-addition

Chlorpromazine hydrochloride (Millipore Sigma) was dissolved in ethanol and ammonium chloride (Millipore Sigma) in 1×PBS (Millipore Sigma). C6/36 cells were seeded at a density of 6 × 10^5^ cells/mL in a 24-well plate and were treated in parallel with the inhibitor or the equivalent amount of vehicle. Cells were infected with ZIKV (MOI 0.1) and inhibitors were added at the indicated time points. Each treatment with CPZ or NH_4_Cl was for four hours, after which time the inhibitors were removed and fresh media was added. The cells were kept at 28°C for 48 hours when the supernatant was collected and titrated.

### ZIKV RNA transfection

Infectious viral RNA was isolated from viral particles. Briefly, five 15-cm dishes of Vero cells were inoculated with ZIKV (MOI 0.01) for six days. Supernatant was clarified (1000 × g; 3 min) to remove the cell debris, and the virus was then pelleted through a 20% sucrose cushion at 37,000 rpm in an F37L-8x100 rotor in a Sorvall WX ultracentrifuge. Viral particle pellets were lysed in 500 μl TRIzol reagent (Thermo Fisher Scientific) and the RNA was extracted as per manufacturer’s instructions. Viral RNA was quantified using a Take3™ Micro-Volume Plate and BioTek microplate reader. Standard and 20°C-adapted C6/36 cells were plated at a density of 7 × 10^5^ cells/mL in 48-well plates the day before transfection. Infectious RNA (3 μg) was diluted in 100 μl of serum free L-15 media and mixed with Lipofectamine™ 3000 (Invitrogen) transfection reagents following the manufacturer’s instruction. The transfection mixture was equally distributed over three wells on three plates. Each plate was incubated for two hours at its respective temperatures. The media was replaced, and some plates were switched to different temperatures. Supernatant was collected at indicated time points, and viral titers were determined using TCID_50_ method.

### Translation quantification

A translation reporter construct, synthesized by Genewiz, that encodes NanoLuc® luciferase with the 5’- and 3’-UTRs of ZIKV was amplified based on the previously published protocol (78). The DNA was *in vitro* transcribed by using the HiScribe T7 ARCA mRNA kit (NEB) and capped RNA was isolated by using an RNA Clean & Concentrator (Zymo Research) following the manufacturer’s instructions. Standard and 20°C-adapted C6/36 cells were plated at a density of 1 × 10^6^ cells/mL in 96-well plates and were incubated at their respective temperatures for three hours. Capped RNA (3 μg) was mixed with Lipofectamine™ 3000 (Invitrogen) transfection reagents, following the manufacturer’s instruction, and used to transfect twelve wells on each of three 96-well plates. After one-hour incubation at the respective temperatures, one plate was shifted from 28°C to 20°C. At the indicated time points, the media was removed from duplicate wells, and the cells were lysed in lysis buffer and stored at 4°C. Once all time points were collected, the Nano-Glo® substrate (Promega) was added, and the luminescence was measured in a Glomax® Explorer (Promega) plate reader.

### Immunocytochemistry

C6/36 cells were plated at a density of 6 × 10^5^ cells/mL in a 24-well plate containing a glass coverslip and incubated with ZIKV (high MOI, >10). Plates were shifted to the 20°C incubator at the indicated time points and cells were harvested 48 hours following infection. Cells were fixed with 4% paraformaldehyde overnight at 4°C. After fixation, the cells were washed three times in 1×PBS, permeabilized with 0.1% Triton X-100 in 1×PBS for 5 minutes, and blocked with 0.1% Triton X-100 in 1×PBS, 2% FBS for 15 minutes. Cells were incubated with mouse monoclonal anti-dsRNA antibody (rJ2, #MABE1134, EMD Millipore) diluted in a blocking solution (1:60) for one hour, followed by three washes in a blocking solution. Alexa Fluor™ Plus 594-conjugated goat anti-mouse antibody (Invitrogen) was diluted in a blocking solution (1:100) and incubated for 30 minutes while protected from the light. The secondary antibody was removed and 0.1% Hoechst dye in a blocking solution was incubated for 15 minutes in the dark. The cells were washed three times with 1×PBS and coverslips were mounted onto slides with ProLong Gold Antifade Reagent (Molecular Probes). Samples were visualized using an 60x oil immersion lens on a Nikon A1R confocal microscope.

### Statistical analysis

Statistical analyses were performed using Prism 9 for macOS (GraphPad Software Inc.). All data are presented as the mean ± standard error of the mean (SEM) and the number of biological replicates for each experiment is indicated in the figure legends. When calculating significance for viral titers, all data were first log transformed. All multiple comparisons were done using either ordinary one-way of two-way analysis of variance (ANOVA) applying Bonferroni’s correction or Dunnett’s correction when comparing to control or mock. A *p*-value of less than 0.05 was considered to be statistically significant and statistical differences are indicated as follows: * *p* < 0.05, ** *p* < 0.01, *** *p* < 0.001, **** *p* < 0.0001.

## Acknowledgement

We thank the University of Texas Medical Branch Arbovirus Reference Collection for providing the viruses and Dr. Ted Ross for providing C6/36 cells. We would like to thank James Barber at the CVM Cytometry Core Facility for technical support on the confocal microscope and flow cytometer. We gratefully thank the members of the Brindley and Murdock labs for thoughtful comments on the project and manuscript. This research received no specific grant from any funding agency in the public, commercial, or not-for-profit sectors.

## References

1. Brady OJ, Gething PW, Bhatt S, Messina JP, Brownstein JS, Hoen AG, Moyes CL, Farlow AW, Scott TW, Hay SI. 2012. Refining the global spatial limits of dengue virus transmission by evidence-based consensus. PLoS Negl Trop Dis 6:e1760.

2. PAHO. 2018. Pan American Health Organisation. http://www.paho.org/. Accessed January 15.

3. Mlakar J, Korva M, Tul N, Popovic M, Poljsak-Prijatelj M, Mraz J, Kolenc M, Resman Rus K, Vesnaver Vipotnik T, Fabjan Vodusek V, Vizjak A, Pizem J, Petrovec M, Avsic Zupanc T. 2016. Zika virus associated with microcephaly. N Engl J Med 374:951–8.

4. Cao-Lormeau V-M, Blake A, Mons S, Lastère S, Roche C, Vanhomwegen J, Dub T, Baudouin L, Teissier A, Larre P, Vial A-L, Decam C, Choumet V, Halstead SK, Willison HJ, Musset L, Manuguerra J-C, Despres P, Fournier E, Mallet H-P, Musso D, Fontanet A, Neil J, Ghawché F. 2016. Guillain-Barre Syndrome outbreak associated with Zika virus infection in French Polynesia: a case-control study. The Lancet 387:1531–1539.

5. World Health Organization. 2016. WHO statement on the first meeting of the International Health Regulations (2005) (IHR 2005) Emergency Committee on Zika virus and observed increase in neurological disorders and neonatal malformations. http://www.who.int/mediacentre/news/statements/2016/1st-emergency-committee-zika/en/. Accessed July 19.

6. Mordecai EA, Caldwell JM, Grossman MK, Lippi CA, Johnson LR, Neira M, Rohr JR, Ryan SJ, Savage V, Shocket MS, Sippy R, Stewart Ibarra AM, Thomas MB, Villena O. 2019. Thermal biology of mosquito-borne disease. Ecol Lett 22:1690–1708.

7. Winokur OC, Main BJ, Nicholson J, Barker CM. 2020. Impact of temperature on the extrinsic incubation period of Zika virus in Aedes aegypti. PLoS Negl Trop Dis 14:e0008047.

8. Lambrechts L, Paaijmans KP, Fansiri T, Carrington LB, Kramer LD, Thomas MB, Scott TW. 2011. Impact of daily temperature fluctuations on dengue virus transmission by Aedes aegypti. Proc Natl Acad Sci U S A 108:7460–5.

9. Xiao FZ, Zhang Y, Deng YQ, He S, Xie HG, Zhou XN, Yan YS. 2014. The effect of temperature on the extrinsic incubation period and infection rate of dengue virus serotype 2 infection in Aedes albopictus. Arch Virol 159:3053–7.

10. Mbaika S, Lutomiah J, Chepkorir E, Mulwa F, Khayeka-Wandabwa C, Tigoi C, Oyoo-Okoth E, Mutisya J, Ng’ang’a Z, Sang R. 2016. Vector competence of Aedes aegypti in transmitting Chikungunya virus: effects and implications of extrinsic incubation temperature on dissemination and infection rates. Virol J 13:114.

11. Mordecai EA, Cohen JM, Evans MV, Gudapati P, Johnson LR, Lippi CA, Miazgowicz K, Murdock CC, Rohr JR, Ryan SJ, Savage V, Shocket MS, Stewart Ibarra A, Thomas MB, Weikel DP. 2017. Detecting the impact of temperature on transmission of Zika, dengue, and chikungunya using mechanistic models. PLoS Negl Trop Dis 11:e0005568.

12. Bellone R, Failloux AB. 2020. The Role of Temperature in Shaping Mosquito-Borne Viruses Transmission. Front Microbiol 11:584846.

13. Adelman ZN, Anderson MA, Wiley MR, Murreddu MG, Samuel GH, Morazzani EM, Myles KM. 2013. Cooler temperatures destabilize RNA interference and increase susceptibility of disease vector mosquitoes to viral infection. PLoS Negl Trop Dis 7:e2239.

14. Murdock CC, Paaijmans KP, Bell AS, King JG, Hillyer JF, Read AF, Thomas MB. 2012. Complex effects of temperature on mosquito immune function. Proc Biol Sci 279:3357–66.

15. Lepock JR. 2005. How do cells respond to their thermal environment? Int J Hyperthermia 21:681–7.

16. Zhang X, Sheng J, Plevka P, Kuhn RJ, Diamond MS, Rossmann MG. 2013. Dengue structure differs at the temperatures of its human and mosquito hosts. Proc Natl Acad Sci U S A 110:6795–9.

17. Fibriansah G, Ng TS, Kostyuchenko VA, Lee J, Lee S, Wang J, Lok SM. 2013. Structural changes in dengue virus when exposed to a temperature of 37 degrees C. J Virol 87:7585–92.

18. Zhang X, Sun L, Rossmann MG. 2015. Temperature dependent conformational change of dengue virus. Curr Opin Virol 12:109–12.

19. Kudlacek ST, Premkumar L, Metz SW, Tripathy A, Bobkov AA, Payne AM, Graham S, Brackbill JA, Miley MJ, de Silva AM, Kuhlman B. 2018. Physiological temperatures reduce dimerization of dengue and Zika virus recombinant envelope proteins. J Biol Chem 293:8922–8933.

20. Ackermann M, Padmanabhan R. 2001. De novo synthesis of RNA by the dengue virus RNA-dependent RNA polymerase exhibits temperature dependence at the initiation but not elongation phase. J Biol Chem 276:39926–37.

21. Meyer A, Freier M, Schmidt T, Rostowski K, Zwoch J, Lilie H, Behrens SE, Friedrich S. 2020. An RNA Thermometer Activity of the West Nile Virus Genomic 3’-Terminal Stem-Loop Element Modulates Viral Replication Efficiency during Host Switching. Viruses 12.

22. Kostyuchenko VA, Lim EX, Zhang S, Fibriansah G, Ng TS, Ooi JS, Shi J, Lok SM. 2016. Structure of the thermally stable Zika virus. Nature 533:425–8.

23. Xie X, Yang Y, Muruato AE, Zou J, Shan C, Nunes BT, Medeiros DB, Vasconcelos PF, Weaver SC, Rossi SL, Shi PY. 2017. Understanding Zika Virus Stability and Developing a Chimeric Vaccine through Functional Analysis. MBio 8.

24. Slon Campos JL, Marchese S, Rana J, Mossenta M, Poggianella M, Bestagno M, Burrone OR. 2017. Temperature-dependent folding allows stable dimerization of secretory and virus-associated E proteins of Dengue and Zika viruses in mammalian cells. Sci Rep 7:966.

25. Tesla B, Demakovsky LR, Mordecai EA, Ryan SJ, Bonds MH, Ngonghala CN, Brindley MA, Murdock CC. 2018. Temperature drives Zika virus transmission: evidence from empirical and mathematical models. Proc Biol Sci 285.

26. Persaud M, Martinez-Lopez A, Buffone C, Porcelli SA, Diaz-Griffero F. 2018. Infection by Zika viruses requires the transmembrane protein AXL, endocytosis and low pH. Virology 518:301–312.

27. Li M, Zhang D, Li C, Zheng Z, Fu M, Ni F, Liu Y, Du T, Wang H, Griffin GE, Zhang M, Hu Q. 2020. Characterization of Zika Virus Endocytic Pathways in Human Glioblastoma Cells. Front Microbiol 11:242.

28. Ohkuma S, Poole B. 1978. Fluorescence probe measurement of the intralysosomal pH in living cells and the perturbation of pH by various agents. Proc Natl Acad Sci U S A 75:3327–31.

29. Stiasny K, Fritz R, Pangerl K, Heinz FX. 2011. Molecular mechanisms of flavivirus membrane fusion. Amino Acids 41:1159–63.

30. Rinkenberger N, Schoggins JW. 2019. Comparative analysis of viral entry for Asian and African lineages of Zika virus. Virology 533:59–67.

31. Watts DM, Burke DS, Harrison BA, Whitmire RE, Nisalak A. 1987. Effect of temperature on the vector efficiency of Aedes aegypti for dengue 2 virus. Am J Trop Med Hyg 36:143–52.

32. Turell MJ. 1993. Effect of environmental temperature on the vector competence of Aedes taeniorhynchus for Rift Valley fever and Venezuelan equine encephalitis viruses. Am J Trop Med Hyg 49:672–6.

33. Rohani A, Wong YC, Zamre I, Lee HL, Zurainee MN. 2009. The effect of extrinsic incubation temperature on development of dengue serotype 2 and 4 viruses in Aedes aegypti (L.). Southeast Asian J Trop Med Public Health 40:942–50.

34. Zouache K, Fontaine A, Vega-Rua A, Mousson L, Thiberge JM, Lourenco-De-Oliveira R, Caro V, Lambrechts L, Failloux AB. 2014. Three-way interactions between mosquito population, viral strain and temperature underlying chikungunya virus transmission potential. Proc Biol Sci 281.

35. Reisen WK, Fang Y, Martinez VM. 2006. Effects of temperature on the transmission of west nile virus by Culex tarsalis (Diptera: Culicidae). J Med Entomol 43:309–17.

36. Samuel GH, Adelman ZN, Myles KM. 2016. Temperature-dependent effects on the replication and transmission of arthropod-borne viruses in their insect hosts. Curr Opin Insect Sci 16:108–113.

37. Mojica KD, Brussaard CP. 2014. Factors affecting virus dynamics and microbial host-virus interactions in marine environments. FEMS Microbiol Ecol 89:495–515.

38. Pujhari S, Brustolin M, Macias VM, Nissly RH, Nomura M, Kuchipudi SV, Rasgon JL. 2019. Heat shock protein 70 (Hsp70) mediates Zika virus entry, replication, and egress from host cells. Emerging microbes & infections 8:8–16.

39. Chavez-Salinas S, Ceballos-Olvera I, Reyes-Del Valle J, Medina F, Del Angel RM. 2008. Heat shock effect upon dengue virus replication into U937 cells. Virus Res 138:111–8.

40. Chu JJ, Leong PW, Ng ML. 2005. Characterization of plasma membrane-associated proteins from Aedes albopictus mosquito (C6/36) cells that mediate West Nile virus binding and infection. Virology 339:249–60.

41. Li XD, Deng CL, Yuan ZM, Ye HQ, Zhang B. 2020. Different Degrees of 5’-to-3’ DAR Interactions Modulate Zika Virus Genome Cyclization and Host-Specific Replication. J Virol 94.

42. Zachariassen KE. 1991. Hypothermia and cellular physiology. Arctic Med Res 50 Suppl 6:13–7.

43. Quinn PJ. 1988. Effects of temperature on cell membranes. Symp Soc Exp Biol 42:237–58.

44. Lim XN, Shan C, Marzinek JK, Dong H, Ng TS, Ooi JSG, Fibriansah G, Wang J, Verma CS, Bond PJ, Shi PY, Lok SM. 2019. Molecular basis of dengue virus serotype 2 morphological switch from 29 degrees C to 37 degrees C. PLoS Pathog 15:e1007996.

45. Goo L, Dowd KA, Smith AR, Pelc RS, DeMaso CR, Pierson TC. 2016. Zika Virus Is Not Uniquely Stable at Physiological Temperatures Compared to Other Flaviviruses. mBio 7.

46. Pindi C, Chirasani VR, Rahman MH, Ahsan M, Revanasiddappa PD, Senapati S. 2020. Molecular Basis of Differential Stability and Temperature Sensitivity of ZIKA versus Dengue Virus Protein Shells. Sci Rep 10:8411.

47. Rossignol ED, Peters KN, Connor JH, Bullitt E. 2017. Zika virus induced cellular remodelling. Cell Microbiol 19.

48. Cortese M, Goellner S, Acosta EG, Neufeldt CJ, Oleksiuk O, Lampe M, Haselmann U, Funaya C, Schieber N, Ronchi P, Schorb M, Pruunsild P, Schwab Y, Chatel-Chaix L, Ruggieri A, Bartenschlager R. 2017. Ultrastructural Characterization of Zika Virus Replication Factories. Cell Rep 18:2113–2123.

49. de Armas-Rillo L, Valera MS, Marrero-Hernandez S, Valenzuela-Fernandez A. 2016. Membrane dynamics associated with viral infection. Rev Med Virol 26:146–60.

50. Harris JR, Racaniello VR. 2003. Changes in rhinovirus protein 2C allow efficient replication in mouse cells. J Virol 77:4773–80.

51. Harris JR, Racaniello VR. 2005. Amino acid changes in proteins 2B and 3A mediate rhinovirus type 39 growth in mouse cells. J Virol 79:5363–73.

52. Perera R, Riley C, Isaac G, Hopf-Jannasch AS, Moore RJ, Weitz KW, Pasa-Tolic L, Metz TO, Adamec J, Kuhn RJ. 2012. Dengue virus infection perturbs lipid homeostasis in infected mosquito cells. PLoS Pathog 8:e1002584.

53. Reid DW, Campos RK, Child JR, Zheng T, Chan KWK, Bradrick SS, Vasudevan SG, Garcia-Blanco MA, Nicchitta CV. 2018. Dengue Virus Selectively Annexes Endoplasmic Reticulum-Associated Translation Machinery as a Strategy for Co-opting Host Cell Protein Synthesis. J Virol 92.

54. Heaton NS, Perera R, Berger KL, Khadka S, Lacount DJ, Kuhn RJ, Randall G. 2010. Dengue virus nonstructural protein 3 redistributes fatty acid synthase to sites of viral replication and increases cellular fatty acid synthesis. Proc Natl Acad Sci U S A 107:17345–50.

55. Jiménez de Oya N, Blázquez AB, Casas J, Saiz JC, Martín-Acebes MA. 2018. Direct Activation of Adenosine Monophosphate-Activated Protein Kinase (AMPK) by PF-06409577 Inhibits Flavivirus Infection through Modification of Host Cell Lipid Metabolism. Antimicrob Agents Chemother 62.

56. Willard KA, Demakovsky L, Tesla B, Goodfellow FT, Stice SL, Murdock CC, Brindley MA. 2017. Zika Virus Exhibits Lineage-Specific Phenotypes in Cell Culture, in Aedes aegypti Mosquitoes, and in an Embryo Model. Viruses 9.

57. Goodfellow FT, Willard KA, Wu X, Scoville S, Stice SL, Brindley MA. 2018. Strain-Dependent Consequences of Zika Virus Infection and Differential Impact on Neural Development. Viruses 10.

58. Anfasa F, Siegers JY, van der Kroeg M, Mumtaz N, Stalin Raj V, de Vrij FMS, Widagdo W, Gabriel G, Salinas S, Simonin Y, Reusken C, Kushner SA, Koopmans MPG, Haagmans B, Martina BEE, van Riel D. 2017. Phenotypic Differences between Asian and African Lineage Zika Viruses in Human Neural Progenitor Cells. mSphere 2.

59. Sheridan MA, Balaraman V, Schust DJ, Ezashi T, Roberts RM, Franz AWE. 2018. African and Asian strains of Zika virus differ in their ability to infect and lyse primitive human placental trophoblast. PLoS One 13:e0200086.

60. Shao Q, Herrlinger S, Zhu YN, Yang M, Goodfellow F, Stice SL, Qi XP, Brindley MA, Chen JF. 2017. The African Zika virus MR-766 is more virulent and causes more severe brain damage than current Asian lineage and dengue virus. Development 144:4114–4124.

61. Udenze D, Trus I, Berube N, Gerdts V, Karniychuk U. 2019. The African strain of Zika virus causes more severe in utero infection than Asian strain in a porcine fetal transmission model. Emerg Microbes Infect 8:1098–1107.

62. Dowall SD, Graham VA, Rayner E, Hunter L, Atkinson B, Pearson G, Dennis M, Hewson R. 2017. Lineage-dependent differences in the disease progression of Zika virus infection in type-I interferon receptor knockout (A129) mice. PLoS Negl Trop Dis 11:e0005704.

63. Wimalasiri-Yapa B, Stassen L, Hu W, Yakob L, McGraw EA, Pyke AT, Jansen CC, Devine GJ, Frentiu FD. 2019. Chikungunya Virus Transmission at Low Temperature by Aedes albopictus Mosquitoes. Pathogens 8.

64. Wimalasiri-Yapa B, Barrero RA, Stassen L, Hafner LM, McGraw EA, Pyke AT, Jansen CC, Suhrbier A, Yakob L, Hu W, Devine GJ, Frentiu FD. 2021. Temperature modulates immune gene expression in mosquitoes during arbovirus infection. Open Biol 11:200246.

65. Merwaiss F, Filomatori CV, Susuki Y, Bardossy ES, Alvarez DE, Saleh MC. 2021. Chikungunya Virus Replication Rate Determines the Capacity of Crossing Tissue Barriers in Mosquitoes. J Virol 95.

66. Ferreira PG, Tesla B, Horácio ECA, Nahum LA, Brindley MA, de Oliveira Mendes TA, Murdock CC. 2020. Temperature Dramatically Shapes Mosquito Gene Expression With Consequences for Mosquito-Zika Virus Interactions. Front Microbiol 11:901.

67. Chouin-Carneiro T, David MR, de Bruycker Nogueira F, Dos Santos FB, Lourenço-de-Oliveira R. 2020. Zika virus transmission by Brazilian Aedes aegypti and Aedes albopictus is virus dose and temperature-dependent. PLoS Negl Trop Dis 14:e0008527.

68. Onyango MG, Bialosuknia SM, Payne AF, Mathias N, Kuo L, Vigneron A, DeGennaro M, Ciota AT, Kramer LD. 2020. Increased temperatures reduce the vectorial capacity of Aedes mosquitoes for Zika virus. Emerg Microbes Infect 9:67–77.

69. Blagrove MSC, Caminade C, Diggle PJ, Patterson EI, Sherlock K, Chapman GE, Hesson J, Metelmann S, McCall PJ, Lycett G, Medlock J, Hughes GL, Della Torre A, Baylis M. 2020. Potential for Zika virus transmission by mosquitoes in temperate climates. Proc Biol Sci 287:20200119.

70. Murrieta RA, Garcia-Luna S, Murrieta DJ, Halladay G, Young MC, Fauver JR, Gendernalik A, Weger-Lucarelli J, Rückert C, Ebel GD. 2021. Impact of extrinsic incubation temperature on natural selection during Zika virus infection of *Aedes aegypti*. bioRxiv doi:10.1101/2021.03.02.433538:2021.03.02.433538.

71. Guerbois M, Fernandez-Salas I, Azar SR, Danis-Lozano R, Alpuche-Aranda CM, Leal G, Garcia-Malo IR, Diaz-Gonzalez EE, Casas-Martinez M, Rossi SL, Del Río-Galván SL, Sanchez-Casas RM, Roundy CM, Wood TG, Widen SG, Vasilakis N, Weaver SC. 2016. Outbreak of Zika virus infection, Chiapas State, Mexico, 2015, and first confirmed transmission by Aedes aegypti mosquitoes in the Americas. The Journal of Infectious Diseases 214:1349–1356.

72. Faria NR, Azevedo R, Kraemer MUG, Souza R, Cunha MS, Hill SC, Thézé J, Bonsall MB, Bowden TA, Rissanen I, Rocco IM, Nogueira JS, Maeda AY, Vasami F, Macedo FLL, Suzuki A, Rodrigues SG, Cruz ACR, Nunes BT, Medeiros DBA, Rodrigues DSG, Queiroz ALN, da Silva EVP, Henriques DF, da Rosa EST, de Oliveira CS, Martins LC, Vasconcelos HB, Casseb LMN, Simith DB, Messina JP, Abade L, Lourenço J, Alcantara LCJ, de Lima MM, Giovanetti M, Hay SI, de Oliveira RS, Lemos PDS, de Oliveira LF, de Lima CPS, da Silva SP, de Vasconcelos JM, Franco L, Cardoso JF, Vianez-Júnior J, Mir D, Bello G, Delatorre E, Khan K, et al. 2016. Zika virus in the Americas: Early epidemiological and genetic findings. Science 352:345–349.

73. Haddow AD, Schuh AJ, Yasuda CY, Kasper MR, Heang V, Huy R, Guzman H, Tesh RB, Weaver SC. 2012. Genetic characterization of Zika virus strains: geographic expansion of the Asian lineage. PLoS Negl Trop Dis 6:e1477.

74. Lay Mendoza MF, Acciani MD, Levit CN, Santa Maria C, Brindley MA. 2020. Monitoring Viral Entry in Real-Time Using a Luciferase Recombinant Vesicular Stomatitis Virus Producing SARS-CoV-2, EBOV, LASV, CHIKV, and VSV Glycoproteins. Viruses 12.

75. Mainou BA, Zamora PF, Ashbrook AW, Dorset DC, Kim KS, Dermody TS. 2013. Reovirus cell entry requires functional microtubules. mBio 4.

76. Ramakrishnan MA. 2016. Determination of 50% endpoint titer using a simple formula. World J Virol 5:85–6.

77. González M, Martín-Ruíz I, Jiménez S, Pirone L, Barrio R, Sutherland JD. 2011. Generation of stable Drosophila cell lines using multicistronic vectors. Sci Rep 1:75.

78. Schultz MJ, Tan AL, Gray CN, Isern S, Michael SF, Frydman HM, Connor JH. 2018. Wolbachia wStri Blocks Zika Virus Growth at Two Independent Stages of Viral Replication. mBio 9.

